# Unique roles of vaginal *Megasphaera* phylotypes in reproductive health

**DOI:** 10.1101/2020.08.18.246819

**Authors:** Abigail L. Glascock, Nicole R. Jimenez, Sam Boundy, Vishal N. Koparde, J. Paul Brooks, David J. Edwards, Jerome F. Strauss, Kimberly K. Jefferson, Myrna G. Serrano, Gregory A. Buck, Vaginal Microbiome Consortium, Jennifer M. Fettweis

## Abstract

The composition of the human vaginal microbiome has been extensively studied and is known to influence reproductive health. However, the functional roles of individual taxa and their contributions to negative health outcomes have yet to be well characterized. Here, we examine two vaginal bacterial taxa grouped within the genus *Megasphaera* that have been previously associated with bacterial vaginosis (BV) and pregnancy complications. Phylogenetic analyses support the classification of these taxa as two distinct species. These two phylotypes, *Megasphaera* phylotype 1 (MP1) and *Megasphaera* phylotype 2 (MP2), differ in genomic structure and metabolic potential, suggestive of differential roles within the vaginal environment. Further, these vaginal taxa show evidence of genome reduction and changes in DNA base composition, which may be common features of host dependence and/or adaptation to the vaginal environment. In a cohort of 3,870 women, we observed that MP1 has a stronger positive association with bacterial vaginosis whereas MP2 was positively associated with trichomoniasis. MP1, in contrast to MP2 and other common BV-associated organisms, was not significantly excluded in pregnancy. In a cohort of 52 pregnant women, MP1 was both present and transcriptionally active in 75.4% of vaginal samples. Conversely, MP2 was largely absent in the pregnant cohort. This study provides insight into the evolutionary history, genomic potential and predicted functional role of two clinically relevant vaginal microbial taxa.

---

The vaginal microbiome is an important determinant of women’s reproductive health, pregnancy outcomes and neonatal health^1–6^. Optimal vaginal microbial health is typically characterized by dominance of one or more lactic-acid producing species of the genus *Lactobacillus* that function to lower the pH and prohibit the growth of other organisms^7^. A vaginal microbiome depleted of protective vaginal lactobacilli and enriched in diverse anaerobic species is often clinically diagnosed as bacterial vaginosis (BV). BV is the most common vaginal condition worldwide, affecting an estimated 27% of women in North America^8^. This condition has been associated with an increased risk of acquiring sexually transmitted infections (STIs) as well as pregnancy complications including spontaneous preterm birth^9–12^. While associations of vaginal microbial taxa with reproductive health conditions such as BV are well established, the pathophysiological significance of these taxa remains largely unknown. Developing a more comprehensive understanding of how individual taxa contribute to negative health outcomes is essential for understanding the underlying biological mechanisms and for the development of effective therapeutics.

Here, we focus on two vaginal anaerobic taxa, *Megasphaera* phylotype 1 (MP1) and *Megasphaera* phylotype 2 (MP2) and their roles in reproductive health and disease. Both MP1 and MP2 have been previously associated with bacterial vaginosis across multiple cohorts^3, 13–15^. Due to its high specificity for the condition, MP1 has been used in combination with other taxa for molecular diagnosis of BV^14, 16^. MP2 was described by Martin *et al.* to be more prevalent in samples collected from women with trichomoniasis, suggesting the potential for divergent roles of these vaginal *Megasphaera* in disease states^17^. *Megasphaera* species have also been linked to an increased risk for HIV acquisition^18, 19^. Given that MP1 and MP2 have been observed in the urogenital tracts of adolescent males and heterosexual couples, it seems likely that these bacteria can be sexually transmitted^20, 21^.

Vaginal carriage of *Megasphaera* is strongly associated with BV, and pregnant women with BV have an elevated risk for spontaneous preterm birth^22^. The outcomes across antibiotic intervention studies for prevention of preterm birth have been inconsistent, which may be attributed in part to the significant heterogeneity in study design and the choice and timing of therapeutic intervention^22^. It is now clear that there are different subtypes of BV that can be stratified using molecular approaches, and some subtypes of BV may be more tightly linked to preterm birth than others. Even though BV has long been linked to elevated risk for preterm birth, more recent vaginal microbiome studies have identified higher MP1 carriage in women who go on to deliver preterm^12, 23–25^. Interestingly, Mitchell *et al.* observed MP1 in samples collected from the upper genital tract of women undergoing hysterectomy^26^, suggesting that MP1 may be capable of ascending from the vaginal environment into the upper genital tract. Together, these observations suggest that MP1 can colonize the vaginal environment, ascend into the upper genital tract and potentially contribute to PPROM and/or spontaneous preterm birth.

In the current study, we use several approaches to delineate the roles of MP1 and MP2 in reproductive health. These include phylogenetic analyses that probe the evolutionary history of these organisms, genomic characterization that permits assessment of their metabolic potential, and a study to define their individual associations with demographic and clinical measures.

## RESULTS

### Evolutionary history and genomic divergence of MP1 and MP2

Genomes of three representative MP1 isolates and three representative MP2 isolates were analyzed to gain insight into the mechanisms underlying their colonization of the human vaginal environment (Supplementary Table 1). A phylogenetic analysis of 145 orthologous genes using 110 genomes classified to the class Negativicutes revealed that MP1 and MP2 are evolutionarily distinct and separated from the nearest *Megasphaera*/*Anaeroglobus* clade (Fig. 1). Similar results were observed in a phylogenetic analysis using 16S ribosmal RNA (rRNA) genes and the inferred topology was largely reflective of niche adaptation (Fig. 2). In one case, niche-specific separation did not occur; *Megasphaera* sp. BV3C16-1, which was isolated from the human vagina, was grouped with oral taxa. This taxon has been reported in vaginal microbiome^24^, but it has been observed at low abundance and prevalence. For example, the taxon was identified in five of 3,870 vaginal samples (0.12%) at a threshold of 0.01% in a cohort of women enrolled through the Vaginal Human Microbiome Project (VaHMP)^27^.

**Figure 1.**
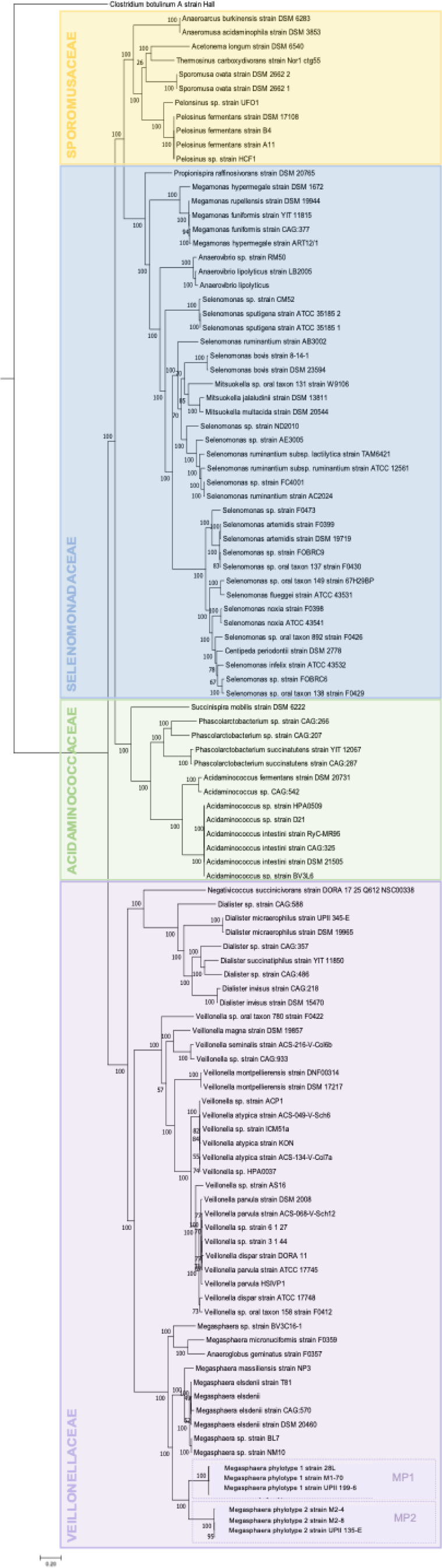
Maximum Likelihood Phylogenetic Tree of the Class Negativicutes. A total of 145 orthologous genes from 110 genomes assigned to the class Negativicutes were included in this analysis. *Clostridium botulinum* A strain Hall was designated as the outgroup. This maximum-likelihood phylogenetic tree was generated using 100 bootstrap replicates. Bootstrap values as present at nodes of the tree. Families within the tree highlighted in different colors: Sporomusaceae: yellow, Selenomonadaceae: blue, Acidaminococcaceae: green, Veillonellaceae: purple. MP1 and MP2 genomes are outlined with dotted lines and labeled.

**Figure 2:**
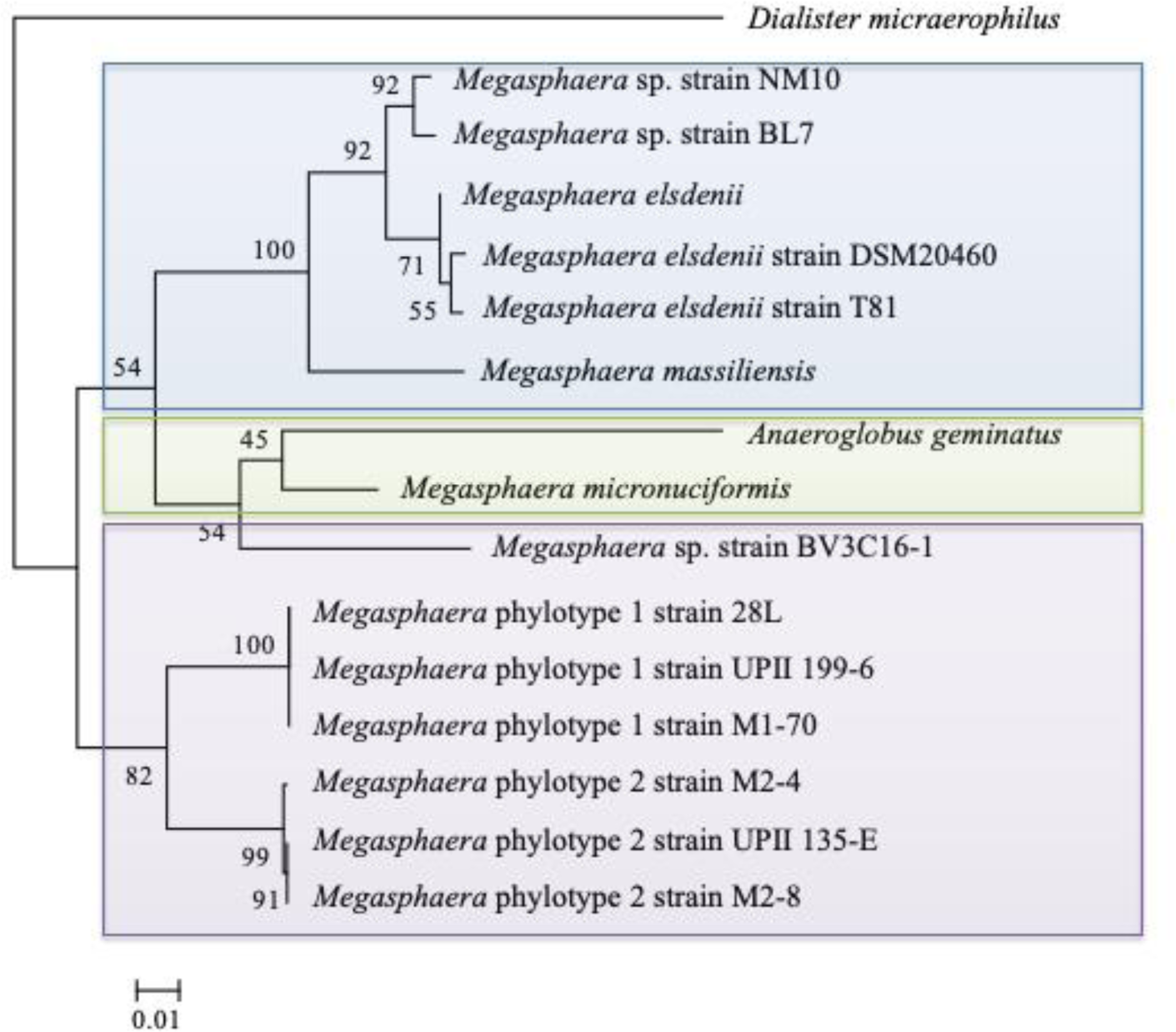
Maximum Likelihood Phylogenetic Tree of 16S Ribosomal RNA gene. This maximum likelihood tree was generated using RAxML-HPC with 1,000 bootstraps. Input data were full-length 16S ribosomal RNA gene sequences (nucleotide). Numbers at nodes are indicative of bootstrap support of that node placement. *Dialister micraerophilus* was selected as the outgroup and is a human oral isolate also classified in the family Veillonellaceae. Remaining isolates are colored by their site of isolation: blue-mammalian gut, green-human oral, purple-human vaginal.

Given the significant divergence of these two vaginal phylotypes from other closely related taxa, we performed a percentage of conserved proteins (POCP) analysis, a metric for delineating genus boundaries^28^. The suggested cutoff for delineation of genera is a POCP value of less than 50% to the genus type strain. The POCP values for members of the MP1 and MP2 clade in comparison to the type strain (*Megasphaera elsdenii* DSM 20460) range from 49.6-52.6% (Supplementary Fig. 1, Supplementary Table 2). *Anaeroglobus geminatus,* currently classified as a separate genus, had a POCP value of 52.5% compared to the *Megasphaera* type strain^29^. A recent study by Campbell *et al.* identified Conserved Signature Indels (CSIs) and Conserved Signature Proteins (CSPs) used to classify organisms to families within the class Negativicutes^30^. We identified all CSIs and CSPs indicative of Veillonellaceae family genomes in MP1 and MP2 genomes (Supplementary Table 3), supporting their previous placement within the Veillonellaceae family. However, three of nine CSP markers specific for the class Negativicutes were absent from all MP1 and MP2 genomes, indicative of genome reduction that is not observed in other host-related *Megasphaera*. While biochemical analyses have yet to be performed, the phylogeny, POCP analysis, loss of CSP markers, and specificity of the clade to the vaginal environment could support placement of these phylotypes into a novel genus of bacteria.

We compared the genomes of MP1 and MP2 with genomes of seven *Megasphaera* isolates from human and mammalian GI tracts, the single human oral *Anaeroglobus* isolate and the vaginal *Megasphaera* sp. BV3C16-1 isolate^29, 31–34^. All of the MP1 and MP2 isolates exhibit evidence of genome reduction with an average genome size of 1.71 megabases (Mb) relative to an average genome size of 2.35 Mb for the other studied *Megasphaera* and *Anaeroglobus* genomes (q=0.001, 95% CI [−0.97,−0.32], Kruskal-Wallis test for differences in genome size with FDR correction). The MP1 and MP2 genomes contain a predicted 1,571 protein-coding genes on average, which is significantly fewer than the number of protein-coding genes for the other studied *Megasphaera* and *Anaeroglobus* genomes, which contained an average of 2,116 genes (q=0.00015, 95% CI [−749,−341], Kruskal-Wallis test for differences in predicted gene count with FDR correction). MP1 and MP2 also exhibit lower average GC composition with an average of 42.6% compared to an average of 51.1% in the other host-associated genomes in the *Megasphaera/Anaeroglobus* clade (q=0.0005, 95% CI [−12.17,−4.75], Kruskal-Wallis test for differences in average GC composition with FDR correction) (Fig. 3a). Reduction in genome size and lower GC percentage has been observed in vaginal strains of other bacterial taxa including *Lactobacillus and Gardnerella*, suggesting reductive evolution may be a common feature of adaptation to the vaginal environment^35, 36^.

**Figure 3.**
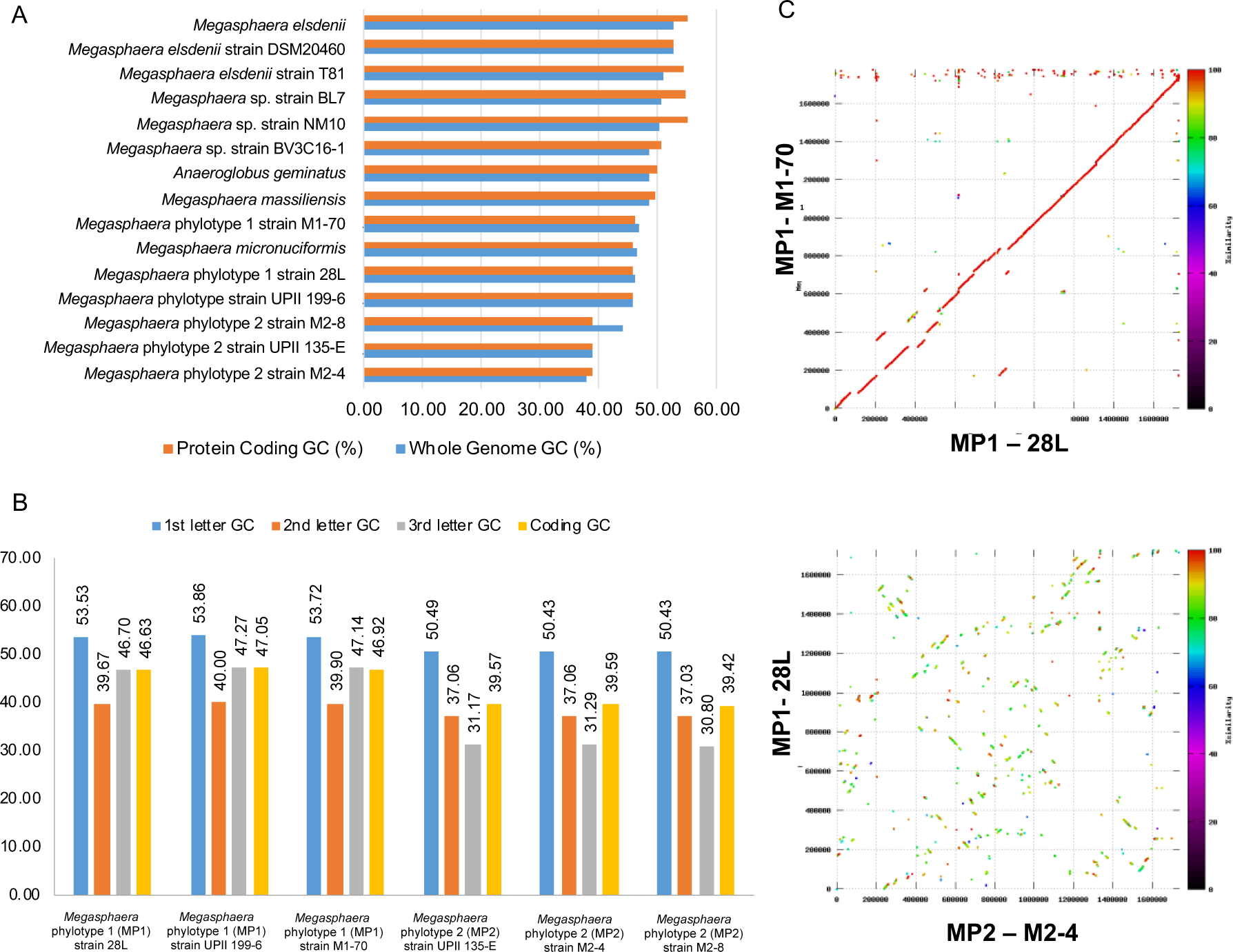
Distinctive GC Composition, Codon Preference & Genomic Structure Between Vaginal *Megasphaera* Phylotypes. a) Differences in both whole genome (blue) and protein-coding (orange) GC composition are shown. b) codon preference is distinct between MP1 and MP2 genomes based on differences in GC composition at specific codon positions (position 1, position 2, position 3 : blue, orange, gray) and in the overall coding GC composition (yellow). c) synteny is conserved within phylotype but lost between MP1 and MP2 genomes. Synteny plots demonstrate structural alignment of genomic content at the amino acid level. Color designates similarity at the amino acid level. The upper panel shows the strong conservation of genomic synteny and protein identity (red color) between two MP1 genomes. The lower panel show massive genome rearrangement and loss of amino acid sequence conservation between a MP1 and a MP2 genome.

### Taxonomic placement of MP1 and MP2 as two discrete species

Similarity of the 16S rRNA gene at an identity threshold of 97% is often used to delineate species. The 16S rRNA similarity between the two phylotypes is 96.3%. This figure along with reports by Srinivasan *et al*., implies that the two phylotypes are best classified as distinct species based on 16S rRNA gene sequence similarity (Supplementary Table 4)^37, 38^. The average nucleotide identity (ANI) between MP1 and MP2, which takes into account the entire nucleotide content of genomes, is 73%. This figure is markedly less than the 95-96% threshold suggested for species demarcation using this method (Supplementary Table 5)^39^. Our phylogenetic analyses (Fig. 1, Fig. 2) reflected these findings, with MP1 and MP2 identified as sister taxa, distinct from other *Megasphaera* and *Anaeroglobus* and separated by significant branch lengths, signifying extensive divergence.

Further comparative analyses revealed that genomic synteny is conserved within phylotype, with variations attributable to the presence of temperate bacteriophage. However, extensive genome rearrangement was observed between MP1 and MP2 genomes (Fig. 3c, Supplementary Fig. 2). While a significant difference in genome size was not observed between MP1 (average of 1.72 Mb) and MP2 (average of 1.70 Mb) isolates (q=0.7497, 95% CI [−0.118, 0.152], Kruskal-Wallis test for differences in genome size with FDR correction), there was an observed difference in GC composition between the MP1 (average of 46.3%) and MP2 isolates (average of 39.0%) (q=0.000002, 95% CI [6.95,7.61], Kruskal-Wallis test for differences in GC composition with FDR correction). The two phylotypes also exhibit GC-divergent codon preference at the third position (average GC composition at third position: MP1-47%, MP2-31%), signaling evolutionary pressure for a reduction in GC composition in MP2 (Fig. 3b). The observed sequence divergence, differential GC composition and codon preference, and lack of synteny between MP1 and MP2 genomes provide support for the designation of the two phylotypes as distinct species.

### Genomic evidence for niche specialization to the vaginal environment

To assess differences in the predicted metabolic potential, we annotated and performed metabolic reconstructions of 15 genomes including representatives of MP1, MP2 and related bacterial strains classified to the *Megasphaera* and *Anaeroglobus* genera. As expected, given the observed genome reduction of MP1 and MP2, many metabolic pathways present among all other related taxa are absent in the MP1/MP2 clade (Supplementary Table 6). MP1 and MP2 are predicted to lack genes conserved in other *Megasphaera* and *Anaeroglobus* genomes that function to transport putrescine and spermidine, metabolize nitrogen, produce selenocysteine and transport and modify the metals nickel and molybdenum. Thus, these organisms may have evolved to rely on synergy with the host and/or microbial co-inhabitants. Interestingly, MP1 and MP2 are predicted to have retained the ability to produce spermidine, a known metabolic marker of BV^40^. Despite the overall genomic reduction of MP1 and MP2, these vaginal phylotypes have also gained functions specific to their clade. MP1 and MP2 specifically encode virulence genes including variable tetracycline resistance genes (*i.e., tetM*, *tetO*, *tetW*) and genes necessary for iron uptake (*i.e., tonB* and hemin uptake outer membrane receptor). Iron sequestration is commonly a critical characteristic of pathogenic bacteria and may be pertinent to the vaginal microbiome given the influx of available iron during menses^41^. MP1 and MP2 genomes also encode multiple CRISPR-associated proteins, which likely function to protect these bacteria from foreign genetic elements^42^.

### Predicted functional divergence of MP1 and MP2

MP1 and MP2 also possess unique predicted metabolic functions, indicative of their divergence. While genomes of both phylotypes encode the majority of genes required for glycolysis, MP2 genomes lack hexokinase. The absence of this gene suggests that MP2 strains cannot use glucose as a carbon source. MP1 genomes are predicted to lack adenosine deaminase (ADA), an enzyme involved in the adenine salvage pathway. In contrast, MP2 genomes retain ADA but lack the gene encoding cytidine deaminase, which functions in the recycling of cytosine bases. These differential salvage strategies are intriguing given that MP1 genomes have markedly higher GC content than MP2 genomes. The phylotypes also differ in their ability to synthesize amino acids. MP2 genomes are incapable of synthesizing leucine and tryptophan, while MP1 genomes lack the ability to interconvert serine and cysteine. Production of aromatic amino acids including tryptophan is energetically expensive^43^. Thus, the loss of tryptophan synthesis genes in MP2 is an example of energetically favorable genome reduction in this host-associated organism.

### MP1 and MP2 phylotypes have distinct clinical associations

Given the distinct metabolic capacities of the MP1 and MP2 phylotypes, we examined the self-survey and clinical data associated with the Vaginal Human Microbiome Project (VaHMP) to investigate their individual roles in reproductive health^27^. We first examined demographic and clinical associations with vaginal MP1 and MP2 carriage in a cohort of 3,091 non-pregnant women. In this cohort, 27% of women (845/3091) carried MP1 only, 5% (163/3091) carried MP2 only, 6% (182/3091) carried both phylotypes, and 62% (1901/3,091) carried neither phylotype. Compared to the average alpha diversity (*i.e.,* inverse Simpson’s index) of samples containing neither of the two phylotypes (1.37), alpha diversity was increased in samples containing MP1 only (1.79), MP2 only (3.47) and both phylotypes (3.37) (Fig. 4). Notably, vaginal microbiome communities containing MP2 exhibited an almost two-fold increase in alpha diversity compared with MP1 alone.

**Figure 4.**
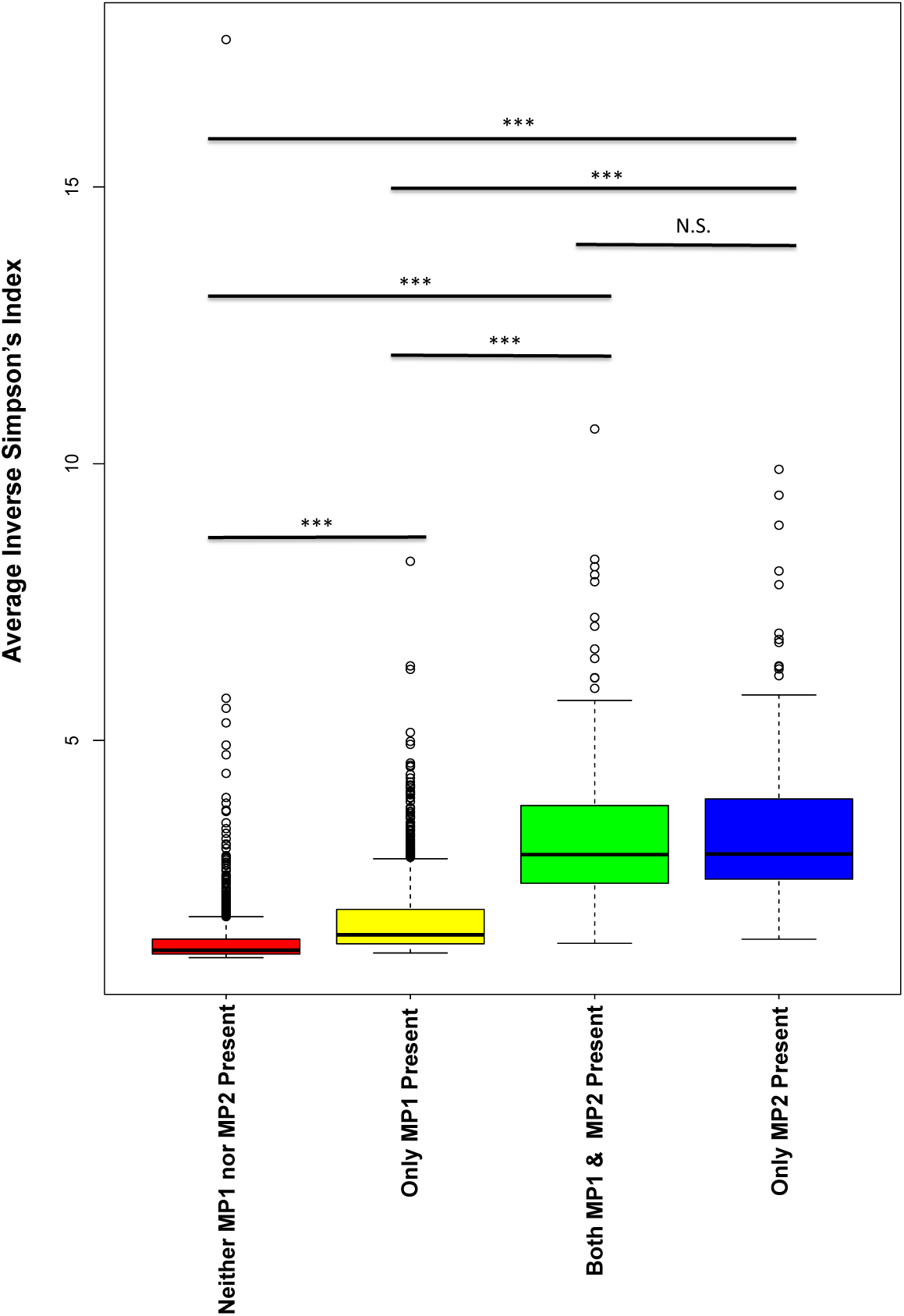
Vaginal *Megasphaera* Phyotypes Associated with Increased Alpha Diversity. Alpha diversity was measured for vaginal microbiome profiles using the Inverse Simpson’s Index, calculated using the ‘vegan’ package in R. Distribution of Inverse Simpson’s Index for each group is shown. Boxes show median and interquartile ranges, with whiskers denoting maximum and minimum values. Outliers are shown as dots. Significance was determined using a two-tailed Student’s T-test. Four different groups are shown, samples containing neither MP1 or MP2 (n=1901, red), samples containing MP1 only (n=845, yellow), samples containing both MP1 and MP2 (n=182, green) and samples containing only MP2 (n=163, blue). Taxa were determined to be present in a sample if they comprised greater than or equal to 0.1% of the sample. Samples with MP1 only, MP2 only and both phylotypes all exhibit increased alpha diveristy, with MP2 only samples being the most highly diverse. All comparisons were found to be highly signifcant (p<0.01) with the exception of MP2 only and both phylotypes.

Associations with demographics were determined using a generalized linear model. Both phylotypes were associated with African-ancestry (MP1: q= 3.00e-31, MP2: q= 1.10e-21, with FDR correction) and a self-reported annual household income of less than 20k (MP1: q= 2.23e-18, MP2: q= 3.31e-18 with FDR correction) (Supplementary Table 7). Fethers *et. al* previously reported that MP1 was associated with women who have sex with women (WSW)^44^. WSW experience higher rates of BV than women who do not have sex with women^45^. Thus, we examined the association of both *Megasphaera* phylotypes with WSW. Although 44% (38/86) of women who reported a current female partner were MP1 positive and there was a positive association between WSW and MP1 carriage (q= 0.075 with FDR correction), it did not reach the threshold for significance (p<0.05). Using the general linearized model (GLM), the race/ethnicity field was identified as a significant covariate with WSW. In stratified analyses, we found that among women who did not report African ancestry, there was a strong association between WSW (N=25) and MP1 (q= 0.0012 with FDR correction), but that among women reporting African ancestry, WSW (N=61) was not significantly associated with MP1 (q= 0.846 with FDR correction). The majority of participants not reporting African ancestry self-identified as Caucasian (68%). This finding highlights the need for precision medicine approaches that account for the contribution of individual environmental and genetic factors and their interactions to fully understand the contributions that shape vaginal microbiome composition and impact risk for adverse reproductive health outcomes.

To assess the association of these two phylotypes with three common vaginal infections (*i.e.,* bacterial vaginosis, candidiasis and trichomoniasis) we performed a relative risk analysis. We observed that while both MP1 and MP2 were associated with an increased risk for BV (MP1: 4.57, 95% CI [3.76,5.55], MP2: 2.19, 95% CI [1.79-2.69]), MP1 is associated with a higher risk for this condition (Table 1). In contrast, MP2 was associated with an increased risk for trichomoniasis (4.84, 95% CI [3.06-7.64]), whereas MP1 had no association (0.96, 95% CI [0.59-1.56]). Using the GLM approach, MP1 and MP2 strains were both associated with self-reported vaginal odor (MP1: q= 5.39e-18, MP2: q= 1.36e-10 with FDR correction) and vaginal discharge (MP1: q= 1.40e-17, MP2: q= 4.64e-7 with FDR correction). Both phylotypes were also associated with clinician-diagnosed elevated vaginal pH (>4.5) (MP1: q= 3.56e-34, MP2: q= 7.29e-12 with FDR correction) consistent with previous reports. Carriage of MP1 and MP2 were also associated with having more than 10 lifetime sexual partners (MP1: q= 0.00037, MP2: q= 4.65e-5 with FDR correction) and having more than one sexual partner in the past month (MP1: q= 0.0002, MP2: q= 2.24e-5 with FDR correction).

**Table 1:** Relative Risk of Vaginal Infections in the Presence of MP1 and MP2

|  | **Bacterial Vaginosis** | **Trichomoniasis** | **Candidiasis** |
| --- | --- | --- | --- |
| ***Megasphaera* phylotype 1** | **4.57 (3.76-5.55)** | **0.96 (0.59-1.56)** | **0.52 (0.37-0.74)** |
| ***Megasphaera* phylotype 2** | **2.19 (1.79-2.69)** | **4.84 (3.06-7.64)** | **0.54 (0.30-0.96)** |
| *Gardnerella vaginalis* | 6.44 (4.18-9.92) | 2.47 (1.23-4.94) | 0.65 (0.49-0.88) |
| *Prevotella* cluster2 | 5.48 (4.31-6.98) | 2.32 (1.44-3.74) | 0.45 (0.33-0.62) |
| Clostridiales BVAB2 | 4.14 (3.46-4.96) | 1.04 (0.63-1.72) | 0.42 (0.28-0.64) |
| *Sneathia amnii* | 3.97 (3.25-4.84) | 2.93 (1.83-4.70) | 0.48 (0.35-0.68) |
| *Mycoplasma hominis* | 2.57 (2.15-3.08) | 5.80 (3.70-9.09) | 1.07 (0.75-1.53) |
| "Ca. *Mycoplasma girerdii*" | 0.88 (0.50-1.55) | 21.00 (13.82-31.91) | 0.71 (0.27-1.87) |

\*Relative risk values were calculated based on the customary relative risk formula for MP1, MP2 and taxa known to be associated with BV and trichomoniasis. Relative Risk = $(A/A+B) / (C/C+D)$ where A represents the number of samples where the taxon is present and the participant is diagnosed with the disease, B represents the number of samples where the taxon is present but the participant is not diagnosed with the disease, C represents the number of samples where the taxon is absent but the participant is diagnosed with the disease and D represents the number of samples where the taxon is not present and the participant is not diagnosed with the disease. The relative risk conferred by each taxon is shown with the 95% confidence interval in parentheses for BV, trichomoniasis and yeast infection (Candidiasis). Our total cohort of non-pregnant women (N=3091) was used for this analysis.

### MP1 and MP2 in Pregnancy

Recent studies have shown that the vaginal microbiome in pregnancy is associated with decreased alpha diversity and dominance of protective *Lactobacillus* species^46–49^. Similarly, BV-associated organisms have been shown to be less prevalent in pregnant women ^27, 50^. Thus, not surprisingly in a case-matched cohort of 779 pregnant and 779 non-pregnant women from the VaHMP study, we found that MP2 was significantly decreased in pregnancy (q< 0.05, Mann-Whitney U test with FDR correction) (Fig. 5). This finding is in agreement with previous work demonstrating that BV organisms are often less prevalent in pregnancy^27, 50^. In contrast, MP1 was not significantly excluded in the pregnant cohort (q= 0.596, Mann-Whitney U test with FDR correction). MP1 has been previously associated with risk for preterm birth^12, 23, 24^; additional studies will be necessary to determine whether the ability of MP1 to persist throughout gestation has implications for complications in pregnancy.

**Figure 5.**
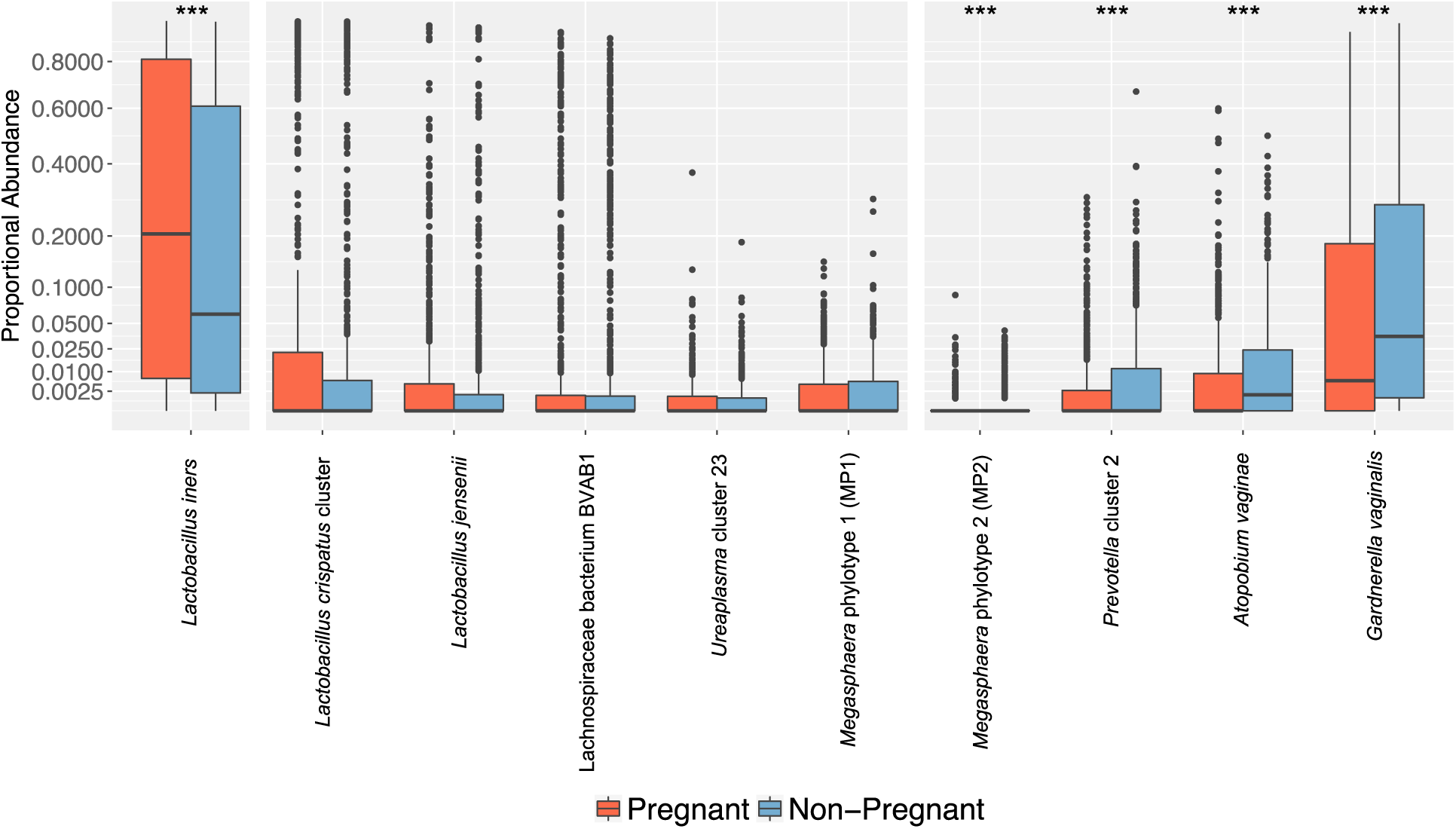
*Megasphaera* phylotype 1 (MP1) Not Significantly Excluded in Pregnancy. Results were generated from a cohort of 779 pregnant women case matched 1:1 with non-pregnant controls (N=1558). Using the R packages ‘wilcox’, a Mann-Whitney U test was performed on all vaginal microbial taxa both present in at least 5% of samples and comprising at least 0.1% relative proportion of the microbiome profile. The R package ‘p.adjust’ was utilized to correct for multiple testing using the FDR correction. The distribution of proportional abundance across both pregnant (red) and non-pregnant (blue) cohorts are shown. Boxes show median and interquartile ranges, with whiskers denoting maximum and minimum values. Outliers are shown as dots. *Lactobacillus iners* is shown to be significantly more prevalent in the pregnant cohort (q=1.20E-6). *Lactobacillus crispatus* cluster*, Lactobacillus jensenii,* Lachnospiraceae BVAB1, *Megasphaera* phylotype 1 (MP1) and *Ureaplasma* cluster 23 are not significantly different between the two cohorts (q=0.19, 0.23, 0.43, 0.56, 0.26 respectively). *Megasphaera* phylotype 2 (MP2), *Prevotella* cluster 2, *Atopobium vaginae* and *Gardnerella vaginalis* are significantly lower in the pregnant cohort (q=5.82×10^−3^, 6.09E-8, 7.95E-8, 2.82E-7 respectively).

To determine whether the two vaginal phylotypes were functionally active in pregnancy, we analyzed metatranscriptomic data from 57 samples collected from 52 pregnant women who delivered at term as a part of the case-control Preterm Birth cohort from the Multi-‘Omic Microbiome Study – Pregnancy Initiative (MOMS-PI)^23^. This is a reanalysis of a subset of an existing dataset previously published in 2019^23, 50^. In this cohort, 43 samples contained transcripts assigned to MP1 while only one sample contained transcripts assigned to MP2 (Supplementary Table 8, Supplementary Table 9), consistent with our observation that MP2 seems to be less prevalent in pregnancy while MP1 is maintained. Because MP2 was only detected in a single sample, we will focus on the findings pertaining to MP1 here. The data showed that *in vivo* in the vaginal environment, MP1 strains transcribed genes from 34 unique pathways. Notably, MP1 strains transcribed genes involved in butyrate production, which has previously been associated with BV^40^.

For this cohort (N=57), we also had paired 16S rDNA profiles and metagenome sequencing profiles generated as a part of a previous study^23^. In these paired data, we observed that the 16S rDNA relative abundance measures for MP1 were strongly correlated to their paired metagenomic relative abundance measures (*ρ*=0.92, Spearman’s rank correlation). This finding supports the use of 16S rDNA profiles in lieu of metagenomic sequencing data to estimate the relative abundance of MP1 in these cohorts. The correlation of MP1 metagenomic relative abundance measures to their paired metatranscriptomic relative abundance measures was also significant (*ρ*=0.91, Spearman’s rank correlation). Intriguingly, the relative abundance measures of the transcripts assigned to MP1 were greater than the observed relative abundance measures in the paired metagenomic dataset (p= 2.95e-05, Mann-Whitney U test) (Fig. 6). This suggests that MP1 is highly transcriptionally active in these samples and makes up a greater proportion of the transcripts than would be predicted based upon the metagenomic data alone. Taken together, the above analyses demonstrate that MP1 is maintained in pregnancy, in contrast to other BV-associated organisms, and is transcriptionally active in a majority of pregnant women in our cohort. These observations in combination with previous associations of MP1 with PPROM and spontaneous preterm labor, highlights MP1 as an important target for future study^12, 23, 24^.

**Figure 6.**
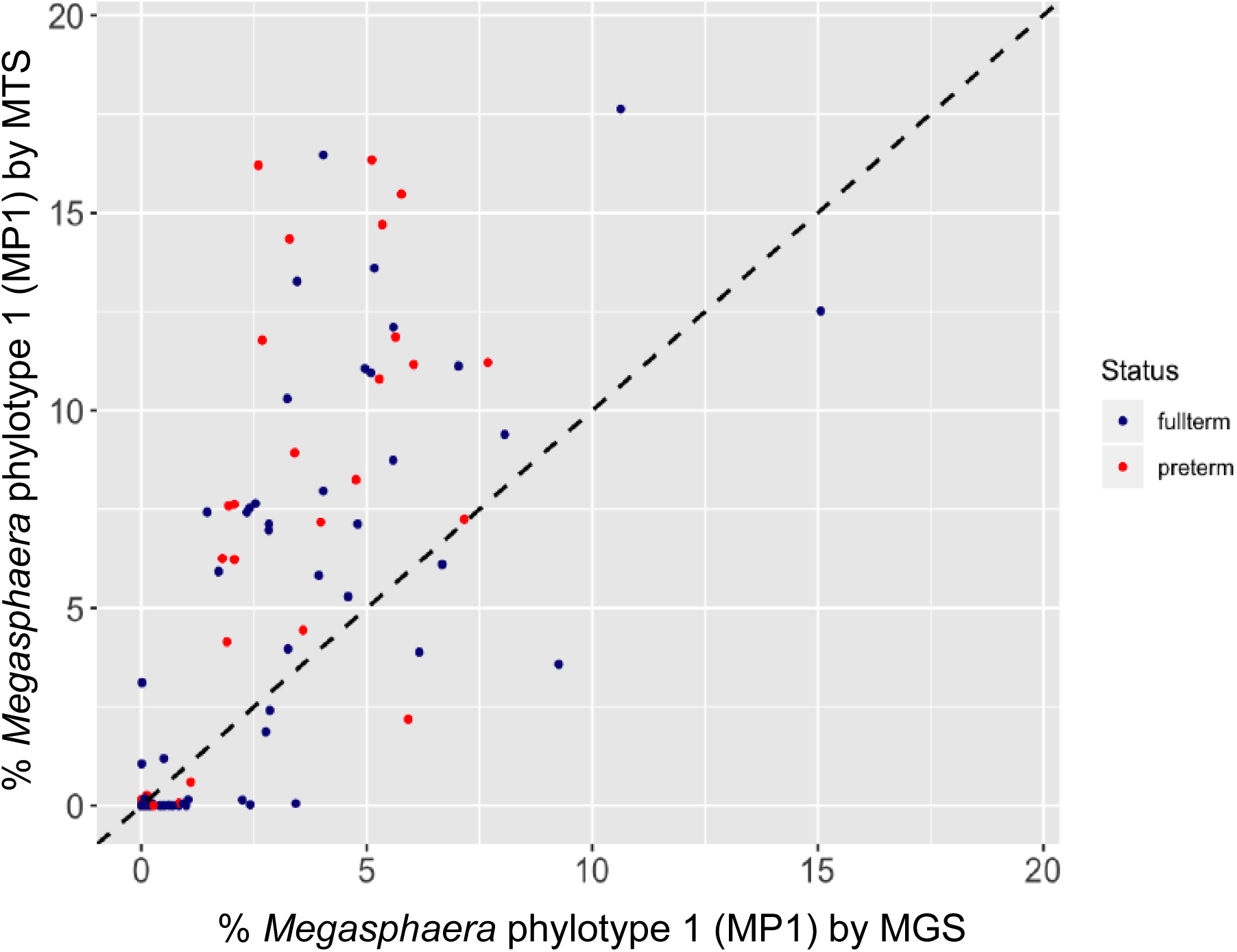
Relationship between *Megasphaera* 16S read abundance and transcript abundance in paired datasets. Results were generated from samples collected from a cohort of pregnant women that participated in the MOMS-PI study. Samples were processed for whole metagenome microbiomics and transcriptomics. Percent of total transcripts attributed to the taxon of interest is shown on the y-axis. Percent of total whole metagenome sequencing reads attributed to the taxon of interest are shown on the x-axis. Each dot represents an individual sample. The relationship between WMGS and WMTS representation of *Megasphaera* phylotype 1 (MP1) is shown. Figures were generated using the R package ‘ggplot’. Data points representing samples from women who went on to deliver full term are shaded blue, while data points representing samples from women who went on to deliver preterm are shaded red. The dotted line extending across the graph diagonally represent the expected 1:1 relationship of WMGS and WMTS-based abundance measures.

## DISCUSSION

In conclusion, our phylogenetic analyses suggest that MP1 and MP2 are evolutionarily divergent from other *Megasphaera* species as well as each other. While comprehensive biological and physiological assays of MP1 and MP2 isolates would be necessary, there is strong phylogenetic evidence that supports placement of MP1 and MP2 into a separate genus. Compared to other *Megasphaera*, both organisms exhibit loss of gut-specific metabolic pathways, acquisition of iron uptake pathways, and loss of genes involved in the biosynthesis of differential amino acids. These organisms also exhibit reduced genomes and lowered GC composition, indicative of a transition to a more host-dependent state^47^ that seems to be a common feature of adaption to the vaginal environment^35, 36^. Taken together these observations are suggestive of adaptation to the host and/or vaginal environment.

Several lines of evidence support the hypothesis that MP1 and MP2 have adapted from an ancestral gastrointestinal tract strain to colonize the vaginal niche: i) the similarity of MP1 and MP2 to human gastrointestinal *Megasphaera* species, ii) the ubiquity of *Megasphaera* in the GI tracts of humans and mammals^31, 32^, iii) the streamlined genomes of MP1 and MP2, a common feature of strains identified in the human vagina, and iv) the physical proximity of the rectum and vagina. Based on our observations, we hypothesize that these two phylotypes share a common ancestor, likely a colonizer of the gastrointestinal tract. Their evolutionary divergence is characterized by progressive gene loss and genome reduction, common features among host-dependent organisms. These changes may be indicative of host dependence and/or adaptation to the vaginal environment specifically.

MP1 and MP2 are evolutionarily divergent and functionally distinct from one another as well, and these findings have important implications for the contributions of these unique phylotypes to vaginal infections and pregnancy complications^3, 12–14, 17, 24^. Several lines of evidence show differential associations of these two phylotypes with clinical diagnoses and demographic factors. As expected, our analyses confirmed that MP1 is tightly correlated with BV as diagnosed by Amsel’s criteria in a cohort of 3,091 non-pregnant women of reproductive age. This result is consistent with numerous previous studies that have demonstrated the strong association of MP1 with BV and led to its use as a biomarker for the diagnosis of the condition8/18/2020 5:45:00 PM. MP2 was also associated with BV (RR= 2.19, 95% C.I. (1.79-2.69)) in the cohort, but to a lesser extent than MP1 (RR= 4.57, 95% C.I. (3.76-5.55)). This finding is consistent with previous reports of the specificity and sensitivity of MP1 and MP2 for BV diagnosis. While MP1 and MP2 have both been reported to have high specificity for BV ranging from 88.5%-98.1% for MP1 and 98.9-100% for MP2 as diagnosed by Amsel’s criteria, Nugent score or a combination of both diagnostic measures, the sensitivity of MP2 (6.9%-31.0%) has been reported to be significantly lower than that of MP1 (68.4%-95.1%)^16, 51^. In the current study, we observed an overall prevalence of 33.2% (n=1027) for MP1 and 11.2% (n=345) for MP2 among non-pregnant women of reproductive age. However, our results also suggest that the two major *Megasphaera* phylotypes may be associated with different subtypes of vaginal dysbiosis.

In the cohort of 3,091 non-pregnant women of reproductive age, MP2 was strongly associated with trichomoniasis whereas MP1 was not associated with the condition. To our knowledge, the association of MP2 with trichomoniasis was first described by Martin *et al.* in 2013, and our study confirms and extends this observation^17^. Martin *et al.* also highlighted an observation from a 1992 study suggesting that *Trichomonas vaginalis* infection was associated with intermediate flora as defined by Nugent score among pregnant women^52^. Together, these findings highlight the need to distinguish between related taxa, such as MP1 and MP2, in microbiome analyses in order to accurately define the functionally relevant subtypes of vaginal dysbiosis and how they contribute to adverse reproductive health outcomes.

In the current study, MP1 and MP2 were both more prevalent among women who reported African ancestry (MP1 AA: 41.1%, MP1 Non-AA: 19.8%, MP2 AA: 15.9%, MP2 Non-AA: 3.1%), consistent with several previous reports including an analysis of the first 1,686 women enrolled in the VaHMP cohort^1, 27, 53^. The association of MP1 and MP2 with African ancestry is consistent with the increased incidence of BV among women with African ancestry^53–55^. In a recent study of 33 white women and 16 black women who were BV negative as assayed by both Amsel’s and Nugent’s criteria, Beamer *et al.* did not detect significant differences in the colonization and density of a number of bacterial species assayed by cultivation and molecular methods^56^. Notably, organisms such as MP1 are rarely observed in women with low Nugent scores; MP1 was identified in 2/33 (6.1%) white women and 2/16 black women (12.5%) by qPCR. While it is less refined than new molecular methods for assaying the vaginal environment, Nugent score, which calculates a score for BV based on the presence of bacterial morphotypes as assayed by microscopy, is still a direct measure of microbial composition. By excluding individuals with higher Nugent score, a significant proportion of women of African ancestry may be excluded from the study. In this current analysis of the VaHMP cohort, race and ethnicity were tightly correlated with a number of covariates including measures of socioeconomic status such as education and annual income. Additional studies will be needed to further define the contributions of both genetic and environmental factors that shape vaginal microbiome composition^55, 57, 58^.

In the current study, MP1 and MP2 were also found to be associated with increased alpha diversity of the microbiome profile and elevated vaginal pH (>4.5), which is one of Amsel’s criteria for BV and is consistent with previous studies linking these *Megasphaera* phylotypes to BV. Interestingly, the vaginal microbiome profile of women who carried MP2 exhibited higher alpha diversity compared to the vaginal microbiome of women who carried MP1 alone. MP2 exhibits greater genome reduction than MP1, likely making it more reliant on other microbial species and/or host factors. This genomic reduction may account for why MP2 is less prevalent in the overall population and specific to a more diverse dysbiotic state.

MP1 is prevalent in the vaginal environment and has been associated with preterm birth in several recent studies, marking it as a taxon of interest^23–25, 49, 50^. Our current study suggests that MP1 levels are similar among pregnant and non-pregnant women, unlike many other BV-associated vaginal taxa which seem to be excluded during the gestational shaping of the vaginal microbiome^25, 59, 60^. MP1 is highly prevalent in the VaHMP cohort, colonizing 33.2% of women in the study. These findings suggest that this highly prevalent organism colonizes the vaginal environment and remains present and transcriptionally active during pregnancy. Mitchell *et al.* observed MP1 in the upper genital tract (UGT) of women undergoing hysterectomy suggesting that this organism is likely capable of ascending into the UGT. This capability combined with the ability of MP1 to maintain colonization during pregnancy suggests that this organism is a candidate for future studies investigating the proposed model where ascending infection of vaginal organisms contributes to in preterm labor and/or birth. *Megasphaera* has also been associated with low vitamin D levels^61^ highlighting a possible link between vaginal microbiome signatures and host state^62^. Identifying mechanisms that permit this organism to pervade the changing vaginal environment associated with the progression of pregnancy may possibly lead to the development of more effective preventative therapeutics targeting microbe-related preterm labor and delivery. This study also highlights the need for continued exploration of mechanisms of microbial evolution in the human microbiome. Understanding the processes that underlie adaptation to specific human host-associated environments will inform strategies for modulating the microbiome to prevent disease and promote human microbial health.

## AUTHOR CONTRIBUTIONS

A.L.G. conducted all experiments and analyses and drafted the manuscript. N.R.J. contributed to clinical association analyses and manuscript preparation. S.B. contributed to comparative genomic analyses. V.N.K. performed genome assembly and initial genome annotation. J.P.B. and D.J.E. provided support for statistical analyses. J.F.S. provided clinically relevant interpretation of results. K.K.J. oversaw cultivation of isolates and provided interpretation of results. M.G.S. oversaw genome sequencing and provided interpretation of phylogenetic data. G.A.B. contributed to interpretation of the results, and G.A.B., K.K.J., J.F.S. and J.M.F. serve as the executive leadership team and planned and directed the overall VaHMP and MOMS-PI studies. The VMC provided infrastructure and data for the study. J.M.F. supervised this study and led the overall direction and planning. A.L.G. and J.M.F designed the study and wrote the manuscript with contributions from all other authors.

## ACKNOWLEDGMENTS

The study team gratefully acknowledges the participants who contributed specimens and data to the Vaginal Human Microbiome Project (VaHMP) and the Multi-Omic Microbiome Project-Pregnancy Initiative (MOMS-PI). The authors would also like to acknowledge other members of the Vaginal Microbiome Consortium and the Research Alliance for Microbiome Science (RAMS) Registry whose contributions made the study possible including the team of research coordinators, the team of sample processors, the team of data managers and the team of clinicians and nurses who assisted with sample collection. We thank our collaborators at the Global Alliance to Prevent Prematurity and Stillbirth (GAPPS) for their contributions to the MOMS-PI study. We acknowledge helpful discussions with Robert P. Hirt, Brian C. Verrelli, Phillip B. Hylemon and Glen E. Kellogg. This study was funded by NIH grants UH3AI083263 and U54HD080784. Other grants that provided partial support include GAPPS BMGF PPB grant and NIH grant R21HD092965. NRJ was supported by grant R25GM090084 for the VCU Initiative For Maximizing Student Development (IMSD) program. All sequence analysis reported herein was performed in the Genomics Core of the Nucleic Acids Research Facilities at VCU; all informatics analysis was performed on servers provided by the Center for High Performance Computing at VCU.

## MATERIALS AND METHODS

### Cultivation of MP1 and MP2

Using anaerobic technique, we cultivated, isolated and sequenced the genomes of one isolate of *Megasphaera* phylotype 1 (MP1, strain M1-70) and two isolates of *Megasphaera* phylotype 2 (MP2, strains M2-4 and M2-8) from frozen glycerol stocks of vaginal swab samples collected through the Vaginal Human Microbiome Project (VaHMP)^63^. One mid-vaginal swab from each participant was used to inoculate 1.0mL of supplemented brain-heart infusion (sBHI) culture media with an added cryo-protectant (20% glycerol) and stored at −80**°**C (Supplementary Table 11). Frozen vaginal culture samples were targeted for cultivation based on the presence and high relative abundance of bacterial targets of interest. These samples were identified using 16S rRNA gene based vaginal microbiome profiles generated for each participant. A scraping of the frozen vaginal culture media from the selected targets was used to inoculate agar plates for bacterial culture. Scrapings were plated on both ThermoScientific Remel Chocolate agar (lysed blood agar) and ThermoScientific Remel Brucella Blood agar (5% sheep’s blood) at four dilutions: 1:10, 1:100, 1:1000 and 1:10000. Plates were stored at 37°C for 24-48 hours. The plates were enclosed in three nested Ziploc bags along with a Mitsubishi Anaeropack-Anaero packet to simulate anaerobic conditions. Individual colonies were selected for growth and purification from the dilution plates based on colony morphology and differential growth characteristics. After re-streaking for visibly pure colonies, the isolates were taxonomically identified by colony PCR amplification of the full 16S rRNA gene using universal 16S primers^64, 65^. Amplicons were purified using the Qiagen QIAquick PCR Purification Kit and sequenced using the Applied Biosystems 3730 DNA Analyzer. Colonies that were identified as bacterial targets of interest and exhibited no evidence of contamination were selected for extraction of genomic DNA. A single colony inoculum was added to 5mL of sBHI in a 15mL falcon tube. Tubes were loosely capped to allow gas exchange and stored in a rack at 37°C for 24-48 hours in three nested Ziploc bags containing a Mitsubishi Anaeropack-Anaero. The DNA was then extracted using the Qiagen DNeasy Blood & Tissue Kit and quantified using the Nanodrop 2000 spectrophotometer. Frozen stocks for MP1 and MP2 isolates were not recoverable.

### Genome Sequencing and Assembly

Purified genomic DNA from the single MP1 isolate was sequenced using the Roche 454 GS FLX Titanium platform. The resulting reads were trimmed for quality and assembled using Newbler v2.8^66^. Purified genomic DNA derived from the two MP2 isolates M2-4 and M2-8 were sequenced using the Illumina MiSeq platform and the resulting reads were trimmed for quality and assembled using Newbler v2.8, CLCBio and SPAdes^66–68^. These three assemblies were merged using CISA to produce the most complete and accurate contigs^69^.

### Structural Genomic Analysis

Genomic synteny was analyzed between genome representatives of MP1 and MP2 and other host-associated *Megasphaera* species. This analysis was performed at the both the protein and nucleic acid level. Nucleic acid-based synteny analyses were performed using NUCmer while amino acid-based synteny analyses were performed using PROmer. Both NUCmer and PROmer are available as a part of the MUMmer 3.0 package^70^. Synteny plots were created using gnuplot from the gnuplot 4.2 package and MUMmerplot, which is also available as a part of the MUMmer 3.0 package^71^. Genomic GC composition was determined using in-house scripts. Codon usage within the genomes was calculated using cusp, a program included in the EMBOSS Tools package available through EMBL-EBI^72^. Comparative analyses of basic genome statistics including genome size, predicted number of proteins and GC composition were performed using a Kruskal-Wallis test. This was performed using the kruskal.test function in R. All calculated p values were adjusted using the FDR correction in R using the p.adjust function. Resulting corrected q values are reported in the Results.

### Measures of Genomic Similarity

Analyses were performed utilizing three MP1 genomes, three MP2 genomes and all publicly available *Megasphaera* and *Anaeroglobus* genomes at NCBI as of January 1, 2015 (Supplementary Table 10). One metagenomic *Megasphaera elsdenii* assembly was excluded from the analysis due to variation in size and gene content from other deposited *M. elsdenii* genomes. One representative of MP1 (Veillonellaceae bacterium DNF00751) and two representatives of MP2 (*Megasphaera* genomosp. 2, Veillonellaceae bacterium KA00182) were deposited after analyses were complete. ANI values suggest that they are similar in genomic content to the genome representatives analyzed in this study. Veillonellaceae bacterium DNF00751 had ANI values ranging from 96.5-98.6% compared to the three MP1 genomes utilized in our analysis. *Megasphaera* genomosp. 2 had ANI values ranging from 98.5-99.0% and Veillonellaceae bacterium KA00182 had ANI values ranging from 98.6-99.0% to the three MP2 genomes utilized in our analyses.

To assess genomic similarity using the entire nucleotide content of the genomes, a pairwise calculation of the average nucleotide identity was performed using a publicly available script (https://github.com/chjp/ANI). 16S ribosomal RNA gene sequences are commonly used to distinguish bacterial species and establish evolutionary relatedness^73^. 16S rRNA gene sequences were identified and extracted from genomes using RNAmmer^74^. Sequence similarity of the 16S rRNA genes was determined using the blastn algorithm^75^. In order to delineate genus boundaries, pairwise Percentage of Conserved Proteins (POCP) values were calculated using in-house scripts developed based on the methods described in Qin et al., 2014^28^.

### CSI and CSP detection

Conserved Signature Proteins (CSPs) and genomic regions containing Conserved Signature Indels (CSIs) were identified using BLAST ^75^. Genomic regions containing CSIs were aligned using MUSCLE and visualized using Jalview^76, 77^. This analysis was based on work performed by Campbell *et al.* identifying CSPs and CSIs indicative of the placement of certain taxa within the class Negativicutes^30^.

### Genome Annotation and Metabolic Reconstruction

Genomes were annotated using both an in-house annotation pipeline and RAST^77^, a web-based genome visualization, annotation and metabolic reconstruction tool provided by NMPDR^78^. As a part of the in-house Genome Annotation Pipeline, the following programs were used. Genes were called using both Glimmer3 and GeneMarkS^79, 80^. Ribosomal RNA genes were identified and extracted from genomes using RNAmmer^74^. Genes encoding tRNAs were identified in genomes using tRNAScan-SE^81^. Orthologous genes were detected using rpsblast in conjunction with Pfam and COG databases^75, 82–85^. Predicted gene functions were annotated using blastx and the nr database at NCBI^75^. Metabolic reconstruction was performed using ASGARD^85^ and visual representations of predicted variation within metabolic pathways were generated using the program color-maps^86^. To determine genes lost in MP1 and MP2, genes specific to MP1 and MP2 and genes that can be used to distinguish the two phylotypes, RAST annotation was utilized. Findings were verified by comparing RAST results to the Genome Annotation Pipeline Glimmer3 and GeneMarkS gene calls. Further verification was performed using the tblastn algorithm to compare known annotated protein sequences available through NCBI to the raw genomic contigs^75, 82, 85^.

### Phylogenetic Analysis

To perform a phylogenetic reconstruction of the 16S rRNA gene, 16S rRNA sequences were identified and extracted from the genomes using RNAmmer^74^. The extracted 16S rRNA gene sequences were aligned using MUSCLE^76^. The resulting alignment file was converted to phylip format using a web-based tool for DNA and protein file format conversion, **AL**ignment **T**ransformation **E**nvi**R**onment or ALTER^87^. RAxML-HPC was used to perform a rapid bootstrap analysis using 1,000 bootstraps and search for the best scoring maximum likelihood tree using the gamma model of heterogeneity^88^. To create a phylogenetic reconstruction of all Negativicutes class genomes, 145 orthologous genes were used. OrthoDB, an online database for orthologous groups was used to determine which orthologous genes were conserved at the family level (Veillonellaceae)^89^. These genes were verified using reciprocal blast and extracted from the six MP1 and MP2 genomes as well as from all publicly available genomes classified to the class Negativicutes at NCBI as of January 1, 2015. *Clostridium botulinum* A strain Hall was selected as the outgroup. This species was chosen due to its classification in the same phylum (Firmicutes) but different class (Clostridia versus Negativicutes) as compared to the Negativicutes genomes. Each orthologous gene was separately aligned using MUSCLE, a program within the EMBOSS Tools package available through EMBL-EBI^72, 76^. Alignments were visually examined and those with large gaps or likely errors were discarded. Sequences from all orthologs were concatenated together to form one large informative sequence. Concatenated sequences were then pruned for informative regions using Gblocks^90^. The resulting sequences were converted from pir to phylip format using the web tool, ALTER^87^. RAxML-HPC was used to perform a rapid bootstrap analysis using 100 bootstraps and search for the best scoring maximum likelihood tree using optimization of substitution rates, the gamma model of heterogeneity and the WAG amino acid substitution matrix ^88^. Aesthetic changes to the tree were made using TreeDyn^91^.

### Participant Recruitment and Informed Consent

We used samples and data from two existing cohorts for this study, The Vaginal Human Microbiome Project (VaHMP) and the Multi-Omic Microbiome Study–Pregnancy Initiative (MOMS-PI), reviewed and approved by the Institutional Review Board at Virginia Commonwealth University (IRB #HM12169, IRB #HM15527). Samples and data are maintained in the Research Alliance for Microbiome Science (RAMS) Registry at Virginia Commonwealth University (IRB #HM15528). The study was performed with compliance to all relevant ethical regulations. Written informed consent was obtained for all participants and parental permission and assent was obtained for participating minors at least 15 years of age.

### Sample Collection, Vaginal Microbiome Profiling and Analysis

Samples collected as part of the Vaginal Human Microbiome Project (VaHMP) at Virginia Commonwealth University were used for this study as previously described^63^. Briefly, mid-vaginal wall swab samples were collected and DNA was extracted from the swabs using the MoBio Powersoil DNA Isolation Kit. DNA samples were randomized to avoid batch effects and the V1-V3 region of the 16S rRNA gene was amplified using polymerase chain reaction (PCR) and universal primers (Supplementary Table 12)^64, 65^. The amplified 16S rDNA fragments were sequenced using the Roche 454 GS FLX Titanium platform. Sequences were classified using both the Ribosomal Database Project (RDP) classifier and the in-house STIRRUPS (<u>S</u>pecies-level <u>T</u>axon Identification of rDNA <u>R</u>eads using a <u>U</u>SEARCH <u>P</u>ipeline <u>S</u>trategy) classifier to achieve species-level classification (version 10-18-17)^92, 93^. Samples that yielded less that 5,000 reads were excluded from analysis.

Taxa were determined to be present if they comprised at least 0.1% of the vaginal microbiome profile of a given sample. Demographics and health history data was self-reported by the participants. Associations were calculated based on the presence or absence of a taxon of interest (threshold of 0.1% of total reads) in combination with given demographic or clinical data. Statistical significance was calculated using a generalized linear model using logistic regression as implemented in the ‘glm’ function in R. All calculated p values were corrected for multiple testing using the FDR correction method. This was performed in R using the p.adjust function. Adjusted q values are reported in the Results.

### Alpha Diversity Measures

16S rDNA-based vaginal microbiome profiles from the Vaginal Human Microbiome Project (VaHMP) outpatient cohort of non-pregnant subjects was used for this analysis (n=3091). Alpha diversity for each microbiome profile was calculated using relative proportion data, renormalized to exclude unclassified reads (below 97% threshold). Inverse Simpson’s Index was used as the measure of alpha diversity. This metric was calculated using the R package ‘vegan.’ Average Inverse Simpson’s Index alpha diversity measures were generated for four subsets of vaginal microbiome data i) samples containing neither phylotype ii) samples containing only MP1 iii) samples containing only MP2 and iv) samples containing both MP1 and MP2. Presence of a phylotype was denoted by a relative abundance of greater than or equal to 0.1% of the vaginal microbiome profile. Statistical significance was calculated using a two-tailed Student’s T-test with a significance level of 0.05.

### Relative Risk

The non-pregnant, outpatient VaHMP cohort was used for this analysis (n=3091). Samples met the threshold of at least 5,000 reads. Vaginal infection status was determined based on clinician diagnosis at time of visit. Relative risk values and their corresponding 95% confidence interval values were calculated based on the standard relative risk formula. Relative Risk = (A/A+B) / (C/C+D) where A represents the number of samples where the taxon is present and the participant is diagnosed with the disease, B represents the number of samples where the taxon is present but the participant is not diagnosed with the disease, C represents the number of samples where the taxon is absent but the participant is diagnosed with the disease and D represents the number of samples where the taxon is not present and the participant is not diagnosed with the disease.

### Pregnancy Analysis

A case-matched cohort was used for this analysis. A cohort of 779 pregnant women was case-matched 1:1 based on ethnicity, age and income to 779 non-pregnant controls. Using the R package ‘wilcox’, we performed a Mann-Whitney U test on all vaginal microbial taxa present in at least 5% of samples that comprise at least 0.1% relative proportion of the microbiome profile. We utilized the R function ‘p.adjust’ to correct for multiple testing using the FDR correction. Results for three *Lactobacillus* species, MP1, MP2 and select associated organisms associated with dysbiosis are shown^94^.

### Transcriptomic Analyses

The Multi-Omic Microbiome Study-Pregnancy Initiative (MOMS-PI) Preterm Birth cohort was utilized for this analysis^95^. This cohort consists of several hundred thousand samples collected from pregnant women throughout and after their pregnancies. For meta-transcriptomics, we collected a mid-vaginal swab from each participant and pre-processed the sample within an hour of collection by inserting the swab into RNAlater® (Qiagen). These swabs were then processed using MoBio Power Microbiome RNA Isolation kit as described by the manufacturer. Total RNA was depleted of human and microbial rRNA using the Epicentre/Illumina Ribo-Zero Magnetic Epidemiology Kit as described by the manufacturer. Enriched messenger RNA was prepared for sequencing by constructing cDNA libraries using the KAPA Biosystems KAPA RNA HyperPrep Kit. Indexed cDNA libraries were pooled in equimolar amounts and sequenced on the Illumina HiSeq 4000 instrument running 4 multiplexed samples per lane with an average yield of ∼100 Gb/lane, sufficient to provide >100X coverage of the expression profiles of the most abundant 15-20 taxa in a sample. Raw sequence data was demultiplexed into sample-specific fastq files using *bcl2fastq* conversion software from Illumina. Adapter residues were trimmed from both 5’ and 3’ end of the reads using Adapter Removal tool v2.1.3. The sequences were trimmed for quality using *meeptools*^96^, retaining reads with minimum read length of 70b and *meep* (maximum expected error) quality score less than 1. Human reads were identified and removed from each sample by aligning the reads to hg19 build of the human genome using the BWA aligner^97^. Transcripts were classified using HUMAnN2^98, 99^ and shortBRED^100^. Transcripts assigned to either MP1 or MP2 were analyzed for this study.

## Data Availability

The genomes of *Megasphaera* phylotype 1 (MP1, strain M1-70), *Megasphaera* phylotype 2 (MP2, strain M2-4) and *Megasphaera* phylotype 2 (MP2, strain M2-8) have been submitted to DDBJ/ENA/GenBank under accession numbers PTJT00000000, PTJU00000000 and PTJV00000000 respectively. The versions described in this paper are versions PTJT01000000, PTJU01000000 and PTJV01000000. Data from the VaHMP has been deposited under dbGAP Study Accession phs000256.v3.p2. Raw metatranscriptomic sequences from the MOMS-PI project are available at NCBI’s controlled-access dbGaP (Study Accession: phs001523.v1.p1). Access to additional fields can be requested through the RAMS Registry (https://ramsregistry.vcu.edu).

## Code availability

Custom code for GC composition and Percentage of Conserved Protein (POCP) calculations is available at https://github.com/Vaginal-Microbiome-Consortium.

---

**Supplementary Table 1.** Genome Characteristics of *Megasphaera* phylotype 1 and *Megasphaera* phylotype 2 Isolates

|  | **M1-70**<br>**(MP1)** | **28L**<br>**(MP1)** | **UPII 199-6**<br>**(MP1)** | **M2-4**<br>**(MP2)** | **M2-8**<br>**(MP2)** | **UPII 135-E**<br>**(MP2)** |
| --- | --- | --- | --- | --- | --- | --- |
| **Isolation Location** | VCU | JCVI | JCVI | VCU | VCU | JCVI |
| **Predicted Genome Size (Mb)** | 1.78 | 1.73 | 1.64 | 1.74 | 1.71 | 1.65 |
| **GC Percentage** | 46.33 | 46.05 | 46.37 | 38.94 | 39.09 | 38.88 |
| **Number of Contigs** | 129 | 34 | 45 | 311 | 328 | 49 |
| **N50 length (bp)** | 179993 | 156177 | 100595 | 102411 | 131070 | 64000 |
| **Number of Contigs @ N50** | 4 | 5 | 7 | 6 | 5 | 8 |
| **Transcriptome Size (Mb)** | 1.55 | 1.55 | 1.46 | 1.46 | 1.41 | 1.44 |
| **Transcriptome/Genome Ratio** | 0.87219 | 0.89894 | 0.88832 | 0.83913 | 0.82962 | 0.87491 |
| **Number of Predicted Genes** | 1647 | 1715 | 1457 | 1591 | 1508 | 1510 |

**Supplementary Table 2. Percentage of Conserved Proteins Among Vaginal *Megasphaera* and Closely Related Genomes.** Percentage of Conserved Proteins (POCP) values were calculated based on the method described by Qin et al. A POCP value of less than 50% is indicative that two genomes should be classified to separate bacterial genera. Pairwise POCP values are denoted by color: dark blue-80-100%, medium blue-60-80%, light blue-50-60%, white-less than 50%, likely isolates from distinct genera.

**Supplementary Table 3.** Conserved Signature Indel and Conserved Signature Protein Analysis of Vaginal *Megasphaera* Phylotypes

|  | *Megasphaera*<br>Phylotype 1 (MP1)<br>strain 28L | *Megasphaera*<br>Phylotype 1 (MP1)<br>strain UPII 199-6 | *Megasphaera*<br>Phylotype 1 (MP1)<br>strain M1-70 | *Megasphaera*<br>Phylotype 2 (MP2)<br>strain UPII 135-E | *Megasphaera*<br>Phylotype 2 (MP2)<br>strain M2-4 | *Megasphaera*<br>Phylotype 2 (MP2)<br>strain M2-8 |
| --- | --- | --- | --- | --- | --- | --- |
| **Conserved Signature Indels** |  |  |  |  |  |  |
| Specific to Class *Negativicutes* |  |  |  |  |  |  |
| 3-Isopropylmalate dehydratase, large subunit<br>(1 aa deletion, position 30-71) | Present | Present | Present | Present | Present | Present |
| DNA-directed RNA polymerase, subunit<br>sigma (1 aa insertion, position 47-73) | Present | Present | Present | Present | Present | Present |
| Specific to Family *Veillonellaceae* |  |  |  |  |  |  |
| GTP disphosphokinase<br>(1 aa deletion, position 441-476) | Present | Present | Present | Present | Present | Present |
| GTP disphosphokinase<br>(1 aa deletion, position 362-403) | Present | Present | Present | Present | Present | Present |
| **Conserved Signature Proteins** |  |  |  |  |  |  |
| Specific to Class *Negativicutes* |  |  |  |  |  |  |
| SELR_02010 | Present | Present | Present | Present | Present | Present |
| SELR_03110 | Present | Present | Present | Present | Present | Present |
| SELR_03270 | Present | Present | Present | Present | Present | Present |
| SELR_05060 | Present | Present | Present | Present | Present | Present |
| SELR_08460 | Present | Present | Present | Present | Present | Present |
| SELR_10260 | Absent | Absent | Absent | Absent | Absent | Absent |
| SELR_10270 | Absent | Absent | Absent | Absent | Absent | Absent |
| SELR_15360 | Absent | Absent | Absent | Absent | Absent | Absent |
| SELR_06480 | Present | Present | Present | Present | Present | Present |
| Specific to Family *Veillonellaceae* |  |  |  |  |  |  |
| MELS_0132 | Present | Present | Present | Present | Present | Present |
| MELS_0206 | Present | Present | Present | Present | Present | Present |
| MELS_0844 | Present | Present | Present | Present | Present | Present |
| MELS_2049 | Present | Present | Present | Present | Present | Present |

\*Presence of Conserved Signature Proteins (CSPs) and genomic regions containing Conserved Signature Indels (CSIs) indicative the class *Negativicutes* and the family *Veillonellaceae* are shown. All CSIs for both the class and family were detected in MP1 and MP2 genomes. All CSPs indicative of the family *Veillonellaceae* were also identified. Absent CSPs (3/9 indicative of the class *Negativicutes*) are denoted in red.

**Supplementary Table 4. 16S Ribosomal RNA Similarity Matrix Among Vaginal *Megasphaera* Phylotypes and Related Genomes.** Pairwise 16S ribosomal RNA similarity was calculated using full length 16S rRNA sequences and the blastn algorithm. Similarity values are denoted by color: red-98-100%, orange-96-98%, yellow-94-96%,green-92-94%, blue-90-92%. The suggested cutoff for delineating species is 97%.

**Supplementary Table 5. Average Nucleotide Identity Analysis Among Vaginal *Megasphaera* Phylotypes and Related Genomes.** Pairwise Average Nucleotide Identity (ANI) was calculated using a publicly availble script (see Methods). ANI values are denoted by color: yellow-greater than 95%, the suggested cutoff for classifying isolates as the same species, green-80-94.99%, blue-less than 80% ANI.

**Supplementary Table 6. Predicted Metabolic Differences between MP1, MP2 and Closely Related Bacterial Taxa.** Sheet 1: Genes that distinguish vaginal Veillonellaceae from closely related species are shown including three sections: genes largely conserved in *Megasphaera* and *Anaeroglobus* but lost in MP1 and MP2, genes specific to oral and vaginal strains, and genes specific to MP1 and/or MP2. Sheet 2: Genes distinguishing MP1 and MP2 genomes are shown in three sections: genes present only in MP1, genes present only in MP2, and genes that are variable between the two phylotypes. Genes present in a specific genome are denoted with an ‘X’.

**Supplementary Table 7. Vaginal *Megasphaera* Phylotypes exhibit Differential Asscociations with Demographics.** General demographics and clinical measures of the non-pregnant, outpatient cohort (n=3091) are shown. Results are separated into five distinct cohorts: i) the overall cohort (n=3091), ii) participants carrying neither MP1 or MP2 (n=1901), iii) participants carrying MP1 only (n=845), iv) participants carrying MP2 only (n=163) and v) participants carrying both phylotypes (n=182). Counts (left) and percentages (right) are shown for each datapoint with the exception of age and sample pH which are shown as averages.

**Supplementary Table 8. *Megasphaera* Phylotype 1 (MP1) Transcription in Pregnancy.** The Multi- ‘Omic Microbiome Study- Pregnancy Initiative (MOMS-PI) cohort was utilized for this analysis. Transcripts were classified using HUMAnN2^98, 99^ and shortBRED^100^ to specific functional pathways. Pathway abundances attributed to MP1 for each sample are shown. Forty-three samples contained MP1 transcripts.

**Supplementary Table 9. *Megasphaera* Phylotype 2 (MP2) Transcription in Pregnancy.** The Multi- ‘Omic Microbiome Study- Pregnancy Initiative (MOMS-PI) cohort was utilized for this analysis. Transcripts were classified using HUMAnN2^98, 99^ and shortBRED^100^ to specific functional pathways. Pathway abundances attributed to MP2 are shown. One sample contained MP2 transcripts.

**Supplementary Table 10. Genomes Utilized in Phylogenetic Reconstruction of the Class Negativicutes.** Basic genome statistics acquired from NCBI are shown. The genomes included were utilized in the creation of the class Negativicutes phylogenetic tree. These include genomes classified to the class Negativicutes and available at NCBI as of January 1, 2015. Genomes were excluded from analysis if they did not contain all 145 orthologous genes needed for the analysis, were low-quality or were significantly different in size or content from other deposited genomes of the same species potentially indicative of a poor asssembly.

**Supplementary Table 11.** Supplemented Brain-Heart Infusion Recipe

| Ingredient | Quantity |
| --- | --- |
| Brain-Heart Infusion Powder (Oxoid) | 9.25g |
| Yeast Extract | 2.50g |
| Gelatin | 2.50g |
| Dextrose | 0.25g |
| Sucrose | 0.25g |
| Deionized Water | 250mL |

**Supplementary Table 12.**
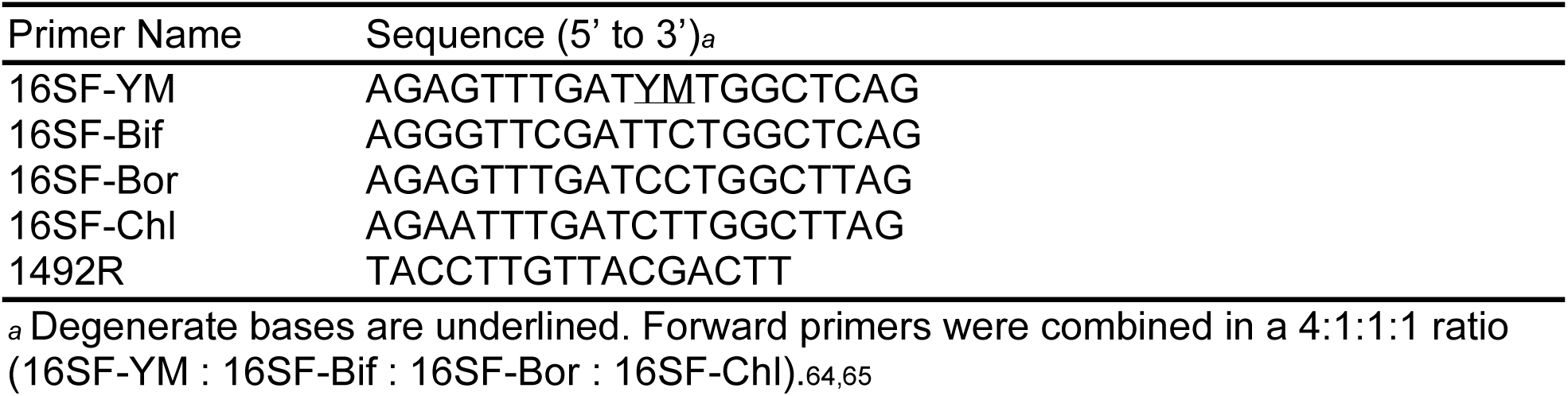
Universal 16S rRNA Gene Primers

**Supplementary Figure 1.**
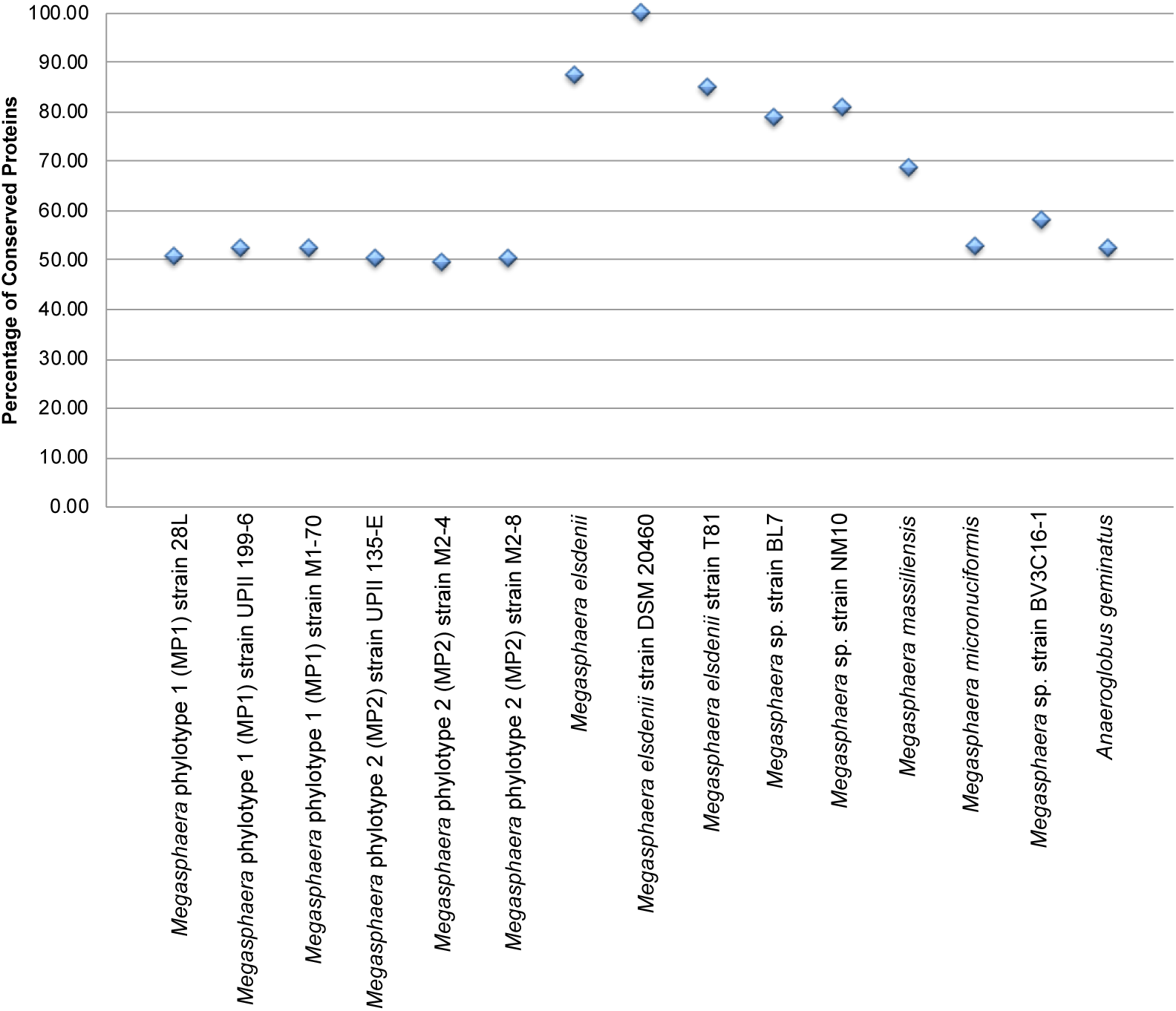
Percentage of Conserved Proteins Analysis versus *Megasphaera* Type Strain. Pairwise Percentage of Conserved Proteins (POCP) values were calculated based on the methods described in Qin et al., 2014. Shown are the POCP values generated between 15 taxa and the *Megasphaera* type strain *Megasphaera elsdenii* strain DSM20460. POCP values below 50% are the suggested cutoff for delineation of a separate bacterial genus.

**Supplementary Figure 2.**
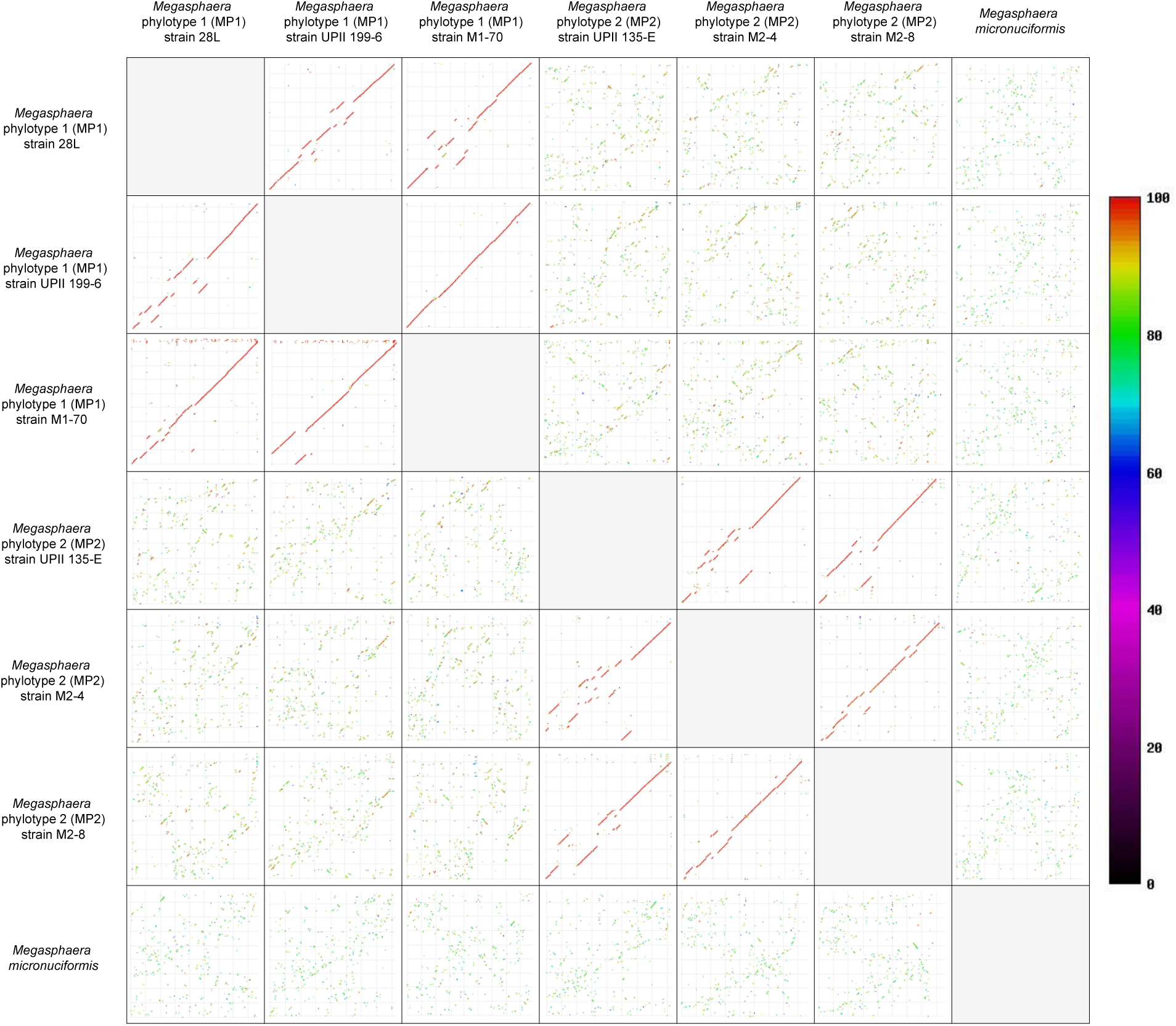
Syntenic Comparison of Vaginal *Megasphaera* Phylotypes and the Oral Isolate *M. micronuciformis*. Full genomes for three MP1, three MP2 and one *Megasphaera micronuciformis* isolate were used for this analysis. Synteny plots demonstrate structural alignment of genomic content at the amino acid level. Color designates similarity at the amino acid level. Synteny is conserved within phylotype as evidenced clear alignment of genomes and protein identity is conserved as well. Between the two phylotypes and in comparsion of *M. micronuciformis*, massive genome rearrangement and loss of amino acid sequence conservation is observed.

